# Assortative mating and mate-choice contributes to a developmental dimorphism in *Streblospio benedicti*

**DOI:** 10.1101/2022.11.27.518082

**Authors:** Erika L. Ruskie, Christina Zakas

## Abstract

Assortative mating, where individuals non-randomly mate with respect to phenotype or genotype, can occur when preferences between potential mates have evolved. When such mate preferences occur in a population it can drive evolutionary and phenotypic divergence. But the extent to which assortative mating, mate preference, and development are evolutionarily linked remains unclear. Here we use *Streblospio benedicti*, a marine annelid with a rare developmental dimorphism, to investigate if mate-choice could contribute to developmental evolution. For *S. benedicti* two types of ecologically and phenotypically similar adults persist in natural populations, but they give rise to distinctly different offspring with alternative life-histories. This dimorphism persists despite the absence of post-zygotic reproductive barriers, where crosses between the developmental types can produce phenotypically intermediate offspring. How this life-history strategy evolved remains unknown, but assortative mating is a typical first step in evolutionary divergence. Here we investigate if female mate-choice is occurring in this species. We find that mate preferences could be contributing to the maintenance of alternative developmental and life-history strategies.

## Introduction

*Streblospio benedicti* is an abundant saltmarsh annelid and one of the best known examples of a phenomenon termed *poecilogony* — where alternative life-history and developmental modes occur within a single species (Giard 1905; Chia et al. 1996; Knott and McHugh 2012). While poecilogony is a rare phenomenon, it receives notable attention in the literature because it is an exception to typical rules of life-history evolution (reviewed in Allen and Pernet 2007). *S. benedicti* is the only known poecilogonous species where developmental traits are heritable, giving rise to two types of females that have numerous reproductive and developmental differences (Levin and Bridges 1994; Gibson et al. 2010; Pernet and McHugh 2010; Zakas and Rockman 2014; Zakas et al. 2018). How these two distinct life-history modes have been maintained in natural populations is a subject of ongoing interest, particularly when considering that crosses of the two developmental types can produce intermediate offspring phenotypes (Levin et al. 1991; Zakas and Rockman 2014, 2021; Zakas et al. 2018). However, these intermediate larval types (and the females that produce them) are rarely, if ever, found in natural populations, suggesting that there is either (or both) strong selection against intermediate phenotypes or that these matings rarely occur (Levin 1984; Schulze et al. 2000; Zakas and Wares 2012). A major evolutionary question is whether mate-choice is contributing to this “non-traditional” life-history strategy and driving population divergence (Collin 2012). Currently it is unclear whether the two persistent developmental types in *S. benedicti* are a case of early incipient speciation, or a stable bet-hedging strategy that rarely evolves. We aim to determine if reproductive isolation arising from mate preferences is contributing to population divergence and developmental evolution in this species.

### *S. benedicti* Mating System

*S. benedicti* is a gonochoristic species where females produce eggs in gametogenic segments throughout their body, and males produce spermatophores that they distribute in the benthos (Fig. 1). Fertilization is internal and embryos move to a transparent dorsal brood pouch where they develop into larvae and are subsequently released (Fig. 1). It is unknown how the females internalize spermatophores, but they can store sperm for at least two months and produce multiple broods. The two developmental strategies of *S. benedicti* are designated by their larval types: Planktotrophic (P) larvae are small, obligately-feeding larvae that develop for two to five weeks in the water column before metamorphosis; Lecithotrophic (L) larvae are about eight times larger, do not require exogenous food, and settle to the benthos within a day of release from the mother’s brood pouch. These alternate life-history strategies and other morphological differences are reviewed in Zakas (2022). Planktotrophy and lecithotrophy are considered evolutionarily stable strategies (ESS) for marine invertebrates that produce larvae. Intermediate larval strategies (such as poecilogony) are rare and presumably selected against (Vance 1973; Hoagland and Robertson 1988; Chia et al. 1996; Allen and Pernet 2007; Zakas and Hall 2012). Despite larval stage differences in ecology and morphology between the two developmental types, adult *S. benedicti* are ecologically and morphologically indistinguishable.

**Figure 1.**
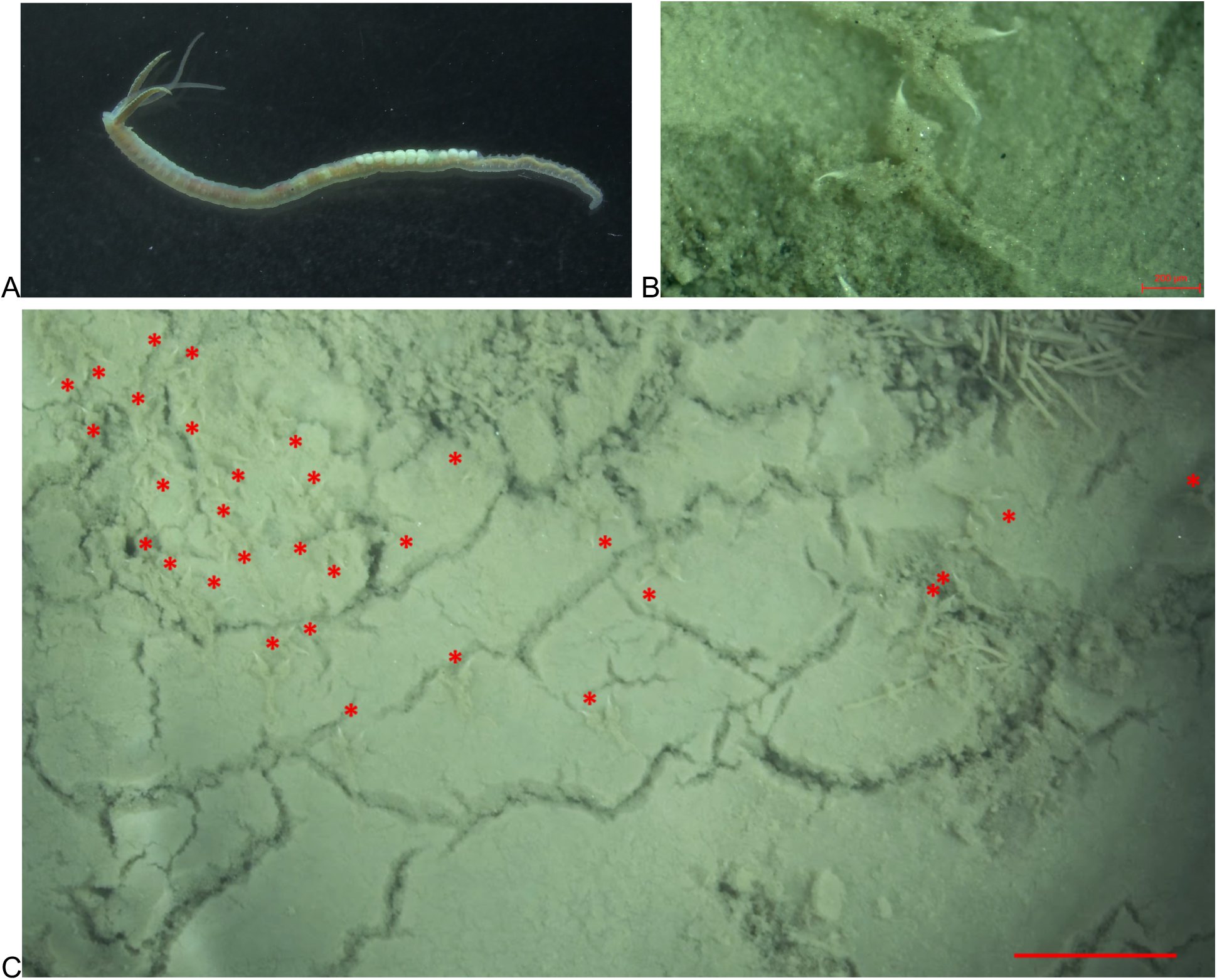
A) Brooding S. benedicti female (~8mm). Paired palps and branchia are the head appendages. B) Each Y-shaped spermatophore has two white sperm tips. Scale bar is 200 μm. C) Males deposit spermatophores throughout the surface of the mud. Red asterisks indicate a spermatophore. Scale bar is 2mm.

Over a decade of no-choice mating crosses in laboratory settings supports that crosses between adults of different developmental types have no noticeable reproductive barriers (Zakas and Rockman 2014, 2021). Other North American populations of different developmental types also confirm the absence of intrinsic reproductive barriers (Levin et al. 1991; Rice 1991; Schulze et al. 2000). Mating is easily achieved by placing males and females together in the same dish. We have observed females selecting spermatophores from the mud using their ciliated palps and branchiae (Fig. 1), but other mating behaviors have not been documented. Laboratory crosses are usually successful provided males are producing spermatophores and females are gravid. Males and females of different developmental types can be crossed in both directions (alternating parental roles), with no noticeable difference in mating frequency, timing, or offspring survival. Offspring production between the two types approximates the frequency within a developmental type or population. Despite highly successful laboratory crosses, females that make intermediate larval types are almost never recovered in natural populations.

### Population Demography

Previous genetic studies demonstrated that populations of *S. benedicti* tend to be predominantly one larval type (Levin 1984; Schulze et al. 2000; Mahon et al. 2009; Zakas and Wares 2012). Although the different populations are sporadically distributed along the US East Coast, they can occur in very close proximity to one another or even overlap. This patchy distribution necessarily leads to assortative mating, where matings occur more often within a group than between groups. Assortative mating is a known premating barrier that can lead to reproductive isolation even in the absence of mate-preferences (Lewontin et al. 1968; Kondrashov and Shpak 1998; Jiang et al. 2013). For *S. benedicti* the encounter rate of mates with the same developmental type is higher, with the exact probability given by the proportion of larval-type alleles in the population (Zakas and Hall 2012). Despite this frequency-based assortative mating, there is evidence of gene flow between neighboring populations. Gene flow on the US East Coast follows a pattern of isolation-by-distance, and geographic proximity acts as a bigger barrier to estimated gene flow than developmental type (Zakas and Wares 2012; Zakas and Rockman 2015). To date, only one larval type has been reported on the West Coast (lecithotrophic individuals have been reported for ~100 years; Carlton 1979) precluding a bicoastal comparison.

In addition to differences in encounter probability, we wanted to identify if other reproductive barriers exist between the developmental types. Identifying barriers to gene flow can indicate where a population stands on a trajectory of divergence and speciation (Nosil 2012; Servedio and Boughman 2017; Kopp et al. 2018). Specifically, female mate preferences favoring males of the same developmental type, and thereby deviating from random mating, could play a major role in the divergence and maintenance of the alternative developmental types (Rosenthal 2017; Kopp et al. 2018). Here we assess whether positive assortative mating arising from female preference for males of the same developmental type (like mates with like) is occurring in *S. benedicti*.

## Methods

### Crosses

Males of two populations: Long Beach, CA (Lecithotrophic, L) and Newark Bay Bayonne, NJ (Planktotrophic, P) were selected when producing spermatophores and paired randomly in a well of a 6-well plate. Pairs of males were kept together for multiple crosses to account for differences in male quality. A female of one developmental type was added to the well, alternating which female (L or P) was introduced first (Fig. 2). Once the female started brooding, she was isolated and the males were moved to a new well, checked for spermatophore production, and a female of the opposite developmental type was added to repeat the experiment. At the end of the experiment, males, females, and offspring were sacrificed for DNA extraction and genotyping. We set up 34 reciprocal crosses (68 total crosses).

**Figure 2.**
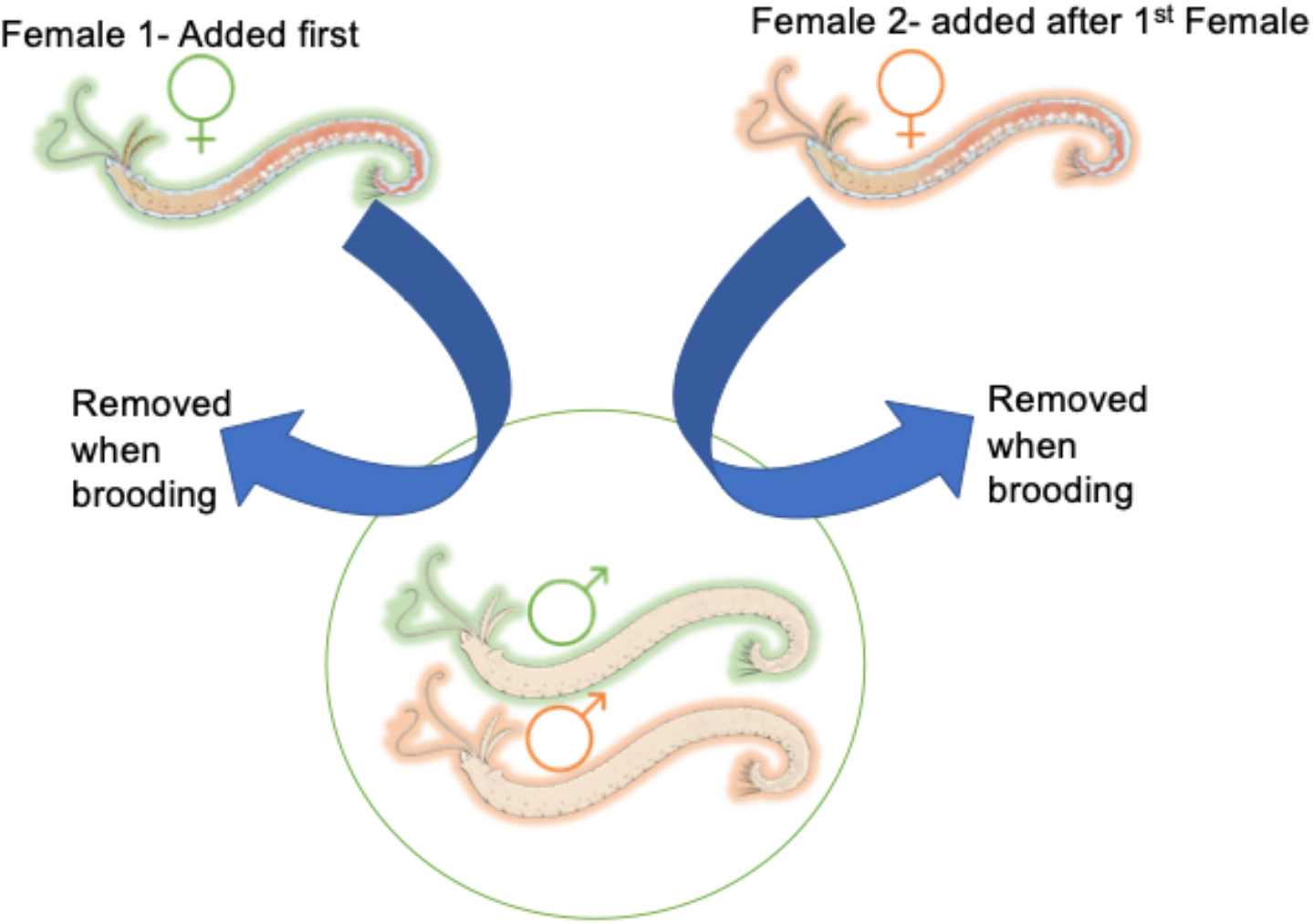
Cross schematic for female mate-choice. The circle indicates a well of a 6-well plate where 2 males of opposite types reside. Green indicates (P) and orange (L). One female is added at a time and removed when she begins brooding offspring.

When females started brooding, they were isolated until they released swimming larvae which were used for DNA extraction (Qiagen DNeasy Blood and Tissue Kit). Eight test crosses had single-paternity clutches (not shown), which indicated we could pool offspring for P mothers. (Subsequent experimental crosses also confirmed single-paternity clutches.) Pooling is necessary as planktotrophic larvae are eight times smaller by volume than lecithotrophic larvae, so obtaining enough DNA from a single planktotrophic larva for PCR was unpredictable. We individually genotyped offspring from L mothers and pooled sets of 10 larvae from P mothers. We genotyped at least eight individual L offspring and at least three pools of P offspring to determine genotypes.

### Genotyping

To determine female mate-choice, we assigned paternity to her offspring. We utilized a single nucleotide polymorphism (SNP) marker that is fixed between the two types. The marker was identified in a previous study by transcriptomic analysis of eight individuals (four of each type; Zakas and Rockman 2015). One of these SNPs, a marker we named SNP_A1, has a thymine fixed in L worms and a cytosine fixed in P worms. This is a cut site for a restriction endonuclease, HinFI, which cuts at a sequence GANTC. SNP_A1 is located on chromosome six at position 1,752,339 in a coding region with uncharacterized gene annotation (Zakas et al. 2022). In addition to the original eight worms from the transcriptome study, we genotyped 50 Ps, 44 Ls, and 20 F_1_s, all of which had the expected genotype at SNP_A1. PCR and restriction digest details are in the supplement.

### Statistical Analysis

We used a binomial exact test in R to determine if females prefer to mate with males of the same type (instances where females chose the same type out of the total trials). We tested for overall female mating preference and within-group female mate preference. We tested for differences in male choice between P and L females using a chi-square test.

## Results

### Genotyping

We successfully genotyped offspring from 53 broods, 28 of which were from L mothers and 25 from P mothers (Fig. 3). We found that 83% of females mated with like-males, which is highly significant (p= 5.6×10^-6^). The proportion was higher for P females, which mated with P males 92% of the time (p=1.6×10^-4^) while L females chose L males 75% of the time (p=0.013).

**Figure 3.**
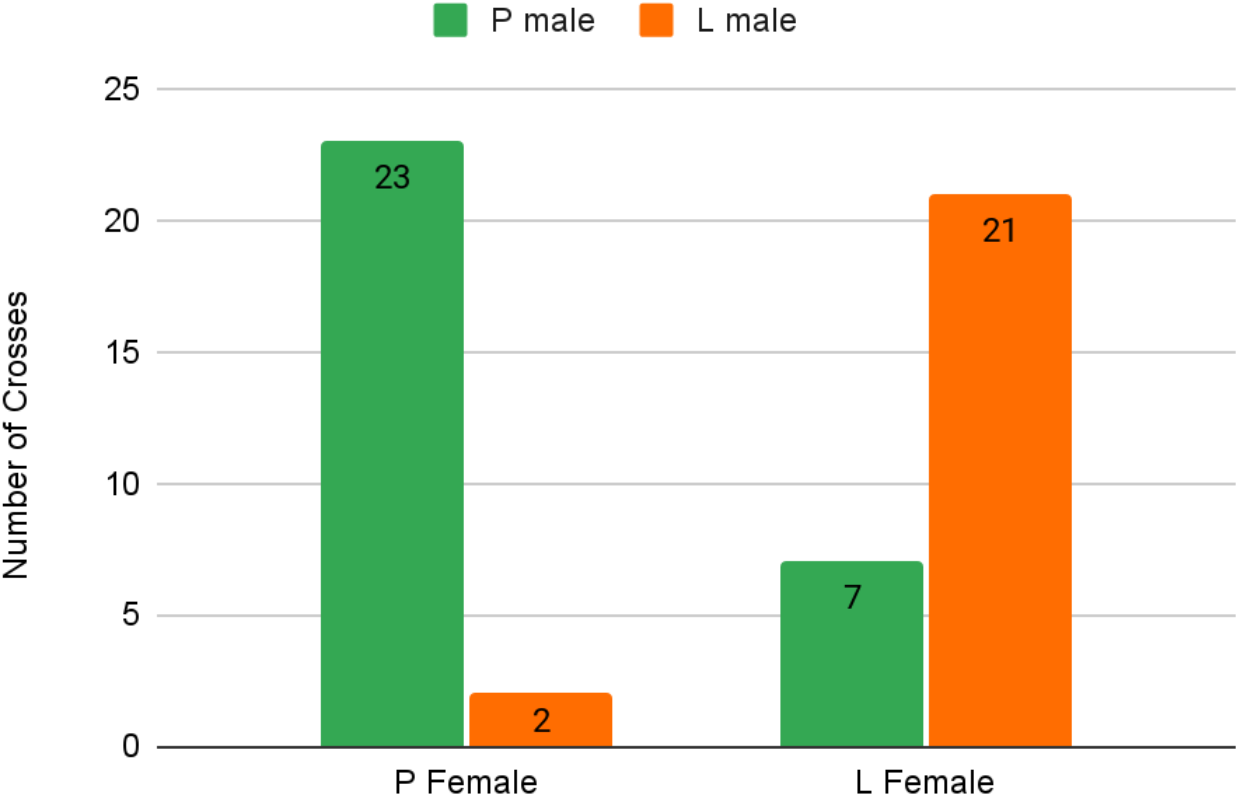
Paternity results for each type of female.

There are 19 sets of crosses where the same pair of males was used with both P and L females. For 13 of the 19 reciprocal crosses, females choose their own type from the pair of males, indicating that there is not a universally “best” male. The order that the females were introduced did not significantly affect the cross outcome (Supplemental Table 1; chi-square test, p=1). For the remaining 15 crosses, one of the males died or stopped producing spermatophores, so the reciprocal cross could not be conducted.

Once we began genotyping animals we noticed that there were a few offspring sets of L mothers that showed a 50:50 ratio of homozygous:heterozygous genotypes indicating that one parent was not conforming to the expected genotypes. Unfortunately, for the crosses in question we were not able to genotype the parents directly. But this led us to genotype other individuals, and we found a few L males in our population were heterozygous. Given all of our prior crosses were single-paternity, we determined this cross result was far more likely to be due to a heterozygous L parent than mixed paternity. This only occurred in three crosses and was revealed because lecithotrophic larvae were genotyped individually. (Further discussion about the potential heterozygosity in these three crosses is in the Supplement.) Removing these three crosses does not change the overall result.

It is important to note that the SNP marker itself is unlikely to be linked to mate-choice, as it is just a marker for paternity assignments. (Although, we do not know what the gene or function of this part of the genome is for *S. benedicti* or how it could be involved in mate selection.) Therefore, we would not assume that the heterozygosity at this marker is affecting female choice.

## Discussion

*S. benedicti* females significantly deviate from random mating and produce offspring with like-males. Although a higher percentage of P females mated with like-males than L females, this overall outcome is not significantly different between the types. While it is clear females are exhibiting some preference (positive assortative mating), we cannot separate the population-of-origin and larval-type of the males used in this study. Therefore we cannot disentangle if female preference is for males of the same developmental type as themselves, or for males from the same population. We plan to investigate if this mate-choice pattern persists in sympatric populations. However, this result indicates that mate-choice can play a major role in maintaining poecilogony in *S. benedicti*.

### Directional patterns of introgression

Our results suggest that more P females mate with same-type males than L females. While this proportion is not significantly different, it could result in directional patterns of introgression in natural populations. Biased directional gene flow could affect the strength of environmental selection on adaptive traits (Harrison 1991; Servedio and Kirkpatrick 1997; Zakas and Hall 2012; de Lafontaine and Bousquet 2017). The degree and direction of female mate preferences will affect the strength of premating barriers, reproductive isolation, and overall population divergence (Feder et al. 2012; Kopp et al. 2018; Richards et al. 2019). Evidence for biased directional gene flow remains to be tested with population data.

Previous studies have also demonstrated phenotypic differences between the F_1_ offspring of alternative crosses (alternating the roles of mother and father: PxL or LxP; Zakas and Rockman 2014). While we have not observed differential survival between the two types of F_1_ offspring, we do see variation in offspring phenotypes, including intermediate trait values. For some traits, F_1_s can phenocopy their mother’s larval type. This is more frequently observed for F_1_s of L mothers. Meaning, it is possible that there is less selection on F_1_s of L mothers given that they can have similar phenotypes to the L offspring (Zakas and Rockman 2014). This could result in weaker mating barriers for L females as their F_1_ offspring may not experience a strong decrease in fitness if these are the larval phenotypes important for survival.

It is currently unknown if F_1_ offspring have decreased fitness in natural populations compared to P or L larvae, but it is reasonable to assume given the prevailing ESS models for marine invertebrates (Vance 1973; Strathmann 1985; Wray and Raff 1991; Levitan 2000). When there is selection on larval fitness it leads to selection for pre-mating isolation barriers (Butlin 1987; Servedio and Kirkpatrick 1997; Nosil 2012, and references therein). This sets the expectation that populations with frequent contact between developmental types (such as a hybrid zone) would have stronger mate-preferences than in homogenous populations (although there are alternatives; Servedio and Boughman 2017). The populations used in this study are from locations that have consistent, homogenous larval types over the ~10 years we have sampled. However, the East Coast population is in close proximity to known lecithotrophic populations (*pers. comm*. Gilligan and Rockman) and gene flow is generally observed across populations of both types on this coast (Zakas and Wares 2012). In contrast, the West Coast lecithotrophs do not encounter individuals of the other developmental type, so we might expect less selection for pre-mating isolation mechanisms to evolve in this population.

### Mate selection mechanism

We do not yet know what behavioral or sensory cues may be important to female matechoice. This could include selection based on male quality, gamete quality, prezygotic “cryptic female choice” or even sperm competition (as females can store sperm). The contribution of these factors remains to be parsed. Premating reproductive isolation strength depends on environmental context and can be ecology-dependent (Endler 1992; Kirkpatrick and Ravigné 2002; Nosil et al. 2005). Lab conditions can alter perception, signaling and mating behavior (Boughman 2002). Further studies are needed to determine the cue(s) for this preference and the strength in natural populations.

One additional limitation to consider is male reproductive behavior. While males were only used if they were visibly making spermatophores at the beginning of the trial, it is possible that they stopped or reduced spermatophore production throughout the experiment. Males will produce spermatophores in isolation, but one study found that males stopped producing spermatophores in the presence of a geographically distant female (Schulze et al. 2000). We attempted to mitigate this factor by only introducing spermatophores to the female in the absence of males, however this never resulted in offspring production (also found in Rice 1991). Here we have attributed positive assortative mating to some aspect of female choice or preference, but the extent that male contribution is involved remains unclear.

### Mate-choice Evolution

Evolved mate-choice preferences have been linked to offspring development mode in some cases (Kelly et al. 2019, 2021*a*, 2021*b*). This potential correlation of developmental mode and mate preference could drive evolutionary shifts in life-history strategy. We might expect this to occur if developmental mode is the target (or correlate) of mate preferences (Servedio and Kopp 2012; Servedio and Boughman 2017). Because selection on sexual traits in natural populations can be stronger than selection on other traits (Kingsolver and Pfennig 2007; Kingsolver and Diamond 2011), such associations could play important roles in the evolution of life-history strategies. The population distribution and poecilogony in *S. benedicti* is an ideal model for investigating these predictions about the interaction of reproductive isolation, lifehistory, and developmental evolution.

## Acknowledgements

Thanks to Nathan Harry and Sarah Cole for feedback and lab assistance. Dhriti Tandon designed and tested the original A1 primers. Kayleigh McHugh provided pictures. Patrick Kelly, Kayleigh McHugh, and Matthew Rockman provided feedback on the manuscript. We received funding from the National Institute of Health: MIRA 5R35GM142853-02 and North Carolina State University startup funds to C. Zakas. E. Ruskie received a Provost’s Professional Experience Program fellowship and an Office of Undergraduate Research Award from NCSU.

## Statement of Authorship

E. Ruskie was an undergraduate honors student in Genetics at NCSU working in the lab of C. Zakas. E. Ruskie was responsible for data collection and analysis as well as writing the original draft. She received an Office of Undergraduate Research Award from NCSU (Summer 2021) to fund the project. C. Zakas conceptualized the methods, acquired additional funding, and wrote the final draft.

## Literature Cited

Allen, J. D., and B. Pernet. 2007. Intermediate modes of larval development: bridging the gap between planktotrophy and lecithotrophy. Evolution & development 9:643–653.

Boughman, J. W. 2002. How sensory drive can promote speciation. Trends in ecology & Evolution 17:571–577.

Butlin, R. 1987. Speciation by reinforcement. Trends in Ecology & Evolution 2:8–13.

Carlton, J. T. 1979. History, biogeography, and ecology of the introduced marine and estuarine invertebrates of the Pacific Coast of North America.

Chia, F. S., G. Gibson, and P. Y. Qian. 1996. Poecilogony as a reproductive strategy of marine invertebrates. Oceanologica Acta 19:203–208.

Collin, R. 2012. Nontraditional life-history choices: what can “intermediates” tell us about evolutionary transitions between modes of invertebrate development? Oxford University Press.

de Lafontaine, G., and J. Bousquet. 2017. Asymmetry matters: A genomic assessment of directional biases in gene flow between hybridizing spruces. Ecology and Evolution 7:3883–3893.

Endler, J. A. 1992. Signals, signal conditions, and the direction of evolution. The American Naturalist 139:S125–S153.

Feder, J. L., S. P. Egan, and P. Nosil. 2012. The genomics of speciation-with-gene-flow. Trends in genetics 28:342–350.

Giard, A. 1905. La poecilogonie. Compte-rendu des Séances du Sixième Congrès International de Zoologie Berne 1904:617–646.

Gibson, G., K. MacDonald, and M. Dufton. 2010. Morphogenesis and phenotypic divergence in two developmental morphs of *Streblospio benedicti* (Annelida, Spionidae). Invertebrate Biology 129:328–343.

Harrison, S. 1991. Local extinction in a metapopulation context: an empirical evaluation. Biological journal of the Linnean Society 42:73–88.

Hoagland, K. E., and R. Robertson. 1988. An assessment of poecilogony in marine invertebrates: phenomenon or fantasy? The Biological Bulletin 174:109–125.

Jiang, Y., D. I. Bolnick, and M. Kirkpatrick. 2013. Assortative mating in animals. The American Naturalist 181:E125–E138.

Kelly, P. W., D. W. Pfennig, S. de la Serna Buzón, and K. S. Pfennig. 2019. Male sexual signal predicts phenotypic plasticity in offspring: implications for the evolution of plasticity and local adaptation. Philosophical Transactions of the Royal Society B 374:20180179.

Kelly, P. W., D. W. Pfennig, and K. S. Pfennig. 2021a. A condition-dependent male sexual signal predicts adaptive predator-induced plasticity in offspring. Behavioral Ecology and Sociobiology 75:1–8.

Kelly, P. W., D. W. Pfennig, and K. S. Pfennig. 2021b. Adaptive Plasticity as a Fitness Benefit of Mate Choice. Trends in Ecology & Evolution 36:294–307.

Kingsolver, J. G., and S. E. Diamond. 2011. Phenotypic selection in natural populations: what limits directional selection? The American Naturalist 177:346–357.

Kingsolver, J. G., and D. W. Pfennig. 2007. Patterns and power of phenotypic selection in nature. Bioscience 57:561–572.

Kirkpatrick, M., and V. Ravigné. 2002. Speciation by natural and sexual selection: models and experiments. The American Naturalist 159:S22–S35.

Knott, K. E., and D. McHugh. 2012. Introduction to symposium: poecilogony—a window on larval evolutionary transitions in marine invertebrates. Integrative and Comparative Biology 52:120–127.

Kondrashov, A. S., and M. Shpak. 1998. On the origin of species by means of assortative mating. Proceedings of the Royal Society of London. Series B: Biological Sciences 265:2273–2278.

Kopp, M., M. R. Servedio, T. C. Mendelson, R. J. Safran, R. L. Rodríguez, M. E. Hauber, E. C. Scordato, et al. 2018. Mechanisms of assortative mating in speciation with gene flow: connecting theory and empirical research. The American Naturalist 191:1–20.

Levin, L. A. 1984. Multiple patterns of development in *Streblospio benedicti* webster (spionidae) from three coasts of North America. The Biological Bulletin 166:494–508.

Levin, L. A., and T. S. Bridges. 1994. Control and consequences of alternative developmental modes in a poecilogonous polychaete. American Zoologist 34:323–332.

Levin, L. A., J. Zhu, and E. Creed. 1991. The genetic basis of life-history characters in a polychaete exhibiting planktotrophy and lecithotrophy. Evolution 45:380–397.

Levitan, D. R. 2000. Optimal egg size in marine invertebrates: theory and phylogenetic analysis of the critical relationship between egg size and development time in echinoids. The American Naturalist 156:175–192.

Lewontin, R., D. Kirk, and J. Crow. 1968. Selective mating, assortative mating, and inbreeding: definitions and implications. Eugenics quarterly 15:141–143.

Mahon, H. K., D. M. Dauer, K. M. Halanych, and A. R. Mahon. 2009. Discrete genetic boundaries of three *Streblospio* (Spionidae, Annelida) species and the status of *S. shrubsolii*. Marine Biology Research 5:172–178.

Nosil, P. 2012. Ecological speciation. Oxford University Press.

Nosil, P., T. H. Vines, and D. J. Funk. 2005. Reproductive isolation caused by natural selection against immigrants from divergent habitats. Evolution 59:705–719.

Pernet, B., and D. McHugh. 2010. Evolutionary changes in the timing of gut morphogenesis in larvae of the marine annelid *Streblospio benedicti*. Evolution & development 12:618–627.

Rice, S. A. 1991. Reproductive isolation in the Polydora ligni complex and the *Streblospio benedicti* complex (Polychaeta: Spionidae). Bulletin of Marine Science 48:432–447.

Richards, E. J., M. R. Servedio, and C. H. Martin. 2019. Searching for sympatric speciation in the genomic era. BioEssays 41:1900047.

Rosenthal, G. G. 2017. Mate choice: the evolution of sexual decision making from microbes to humans. Princeton University Press.

Schulze, S. R., S. A. Rice, J. L. Simon, and S. A. Karl. 2000. Evolution of poecilogony and the biogeography of North American populations of the polychaete *Streblospio*. Evolution 54:1247–1259.

Servedio, M. R., and J. W. Boughman. 2017. The role of sexual selection in local adaptation and speciation. Annu. Rev. Ecol. Evol. Syst 48:85–109.

Servedio, M. R., and M. Kirkpatrick. 1997. The effects of gene flow on reinforcement. Evolution 51:1764–1772.

Servedio, M. R., and M. Kopp. 2012. Sexual selection and magic traits in speciation with gene flow. Current zoology 58:510–516.

Strathmann, R. R. 1985. Feeding and nonfeeding larval development and life-history evolution in marine invertebrates. Annual review of ecology and systematics 16:339–361.

Vance, R. R. 1973. On reproductive strategies in marine benthic invertebrates. The American Naturalist 107:339–352.

Wray, G. A., and R. A. Raff. 1991. The evolution of developmental strategy in marine invertebrates. Trends in Ecology & Evolution 6:45–50.

Zakas, C. 2022. *Streblospio benedicti*: A genetic model for understanding the evolution of development and life-history. Current Topics in Developmental Biology 147:497–521.

Zakas, C., J. M. Deutscher, A. D. Kay, and M. V. Rockman. 2018. Decoupled maternal and zygotic genetic effects shape the evolution of development. eLife 7:e37143.

Zakas, C., and D. W. Hall. 2012. Asymmetric Dispersal Can Maintain Larval Polymorphism: A Model Motivated by *Streblospio benedicti*. Integrative and Comparative Biology 52:197–212.

Zakas, C., N. D. Harry, E. H. Scholl, and M. V. Rockman. 2022. The genome of the poecilogonous annelid *Streblospio benedicti*. Genome biology and evolution 14:evac008.

Zakas, C., and M. V. Rockman. 2014. Dimorphic development in *Streblospio benedicti:* genetic analysis of morphological differences between larval types. International Journal of Developmental Biology 58:593–599.

Zakas, C., and M. V. Rockman. 2015. Gene-based polymorphisms reveal limited genomic divergence in a species with a heritable life-history dimorphism. Evolution & development 17:240–247.

Zakas, C., and M. V. Rockman. 2021. Baby makes three: maternal, paternal, and zygotic genetic effects shape larval phenotypic evolution. Evolution 75:1607–1618.

Zakas, C., and J. P. Wares. 2012. Consequences of a poecilogonous life history for genetic structure in coastal populations of the polychaete *Streblospio benedicti*. Molecular ecology 21:5447–5460.

