## Supplemental Data for "Assortative mating and mate-choice contributes to a developmental dimorphism in *Streblospio benedicti*"

***Supplement Figure 1:*** *Gel electrophoresis designated genotypes. (A) Expected band patterning. (B) Restriction enzyme digestion. Lanes 2-4: single P individuals with homozygous C genotype have a band of 321bp. Lanes 5- 7: Single L individuals with homozygous T genotype and two bands 225 and 96bp. Lanes 8-10: F1 heterozygotes with C/T genotype have all three bands. Lane 11: negative control.*

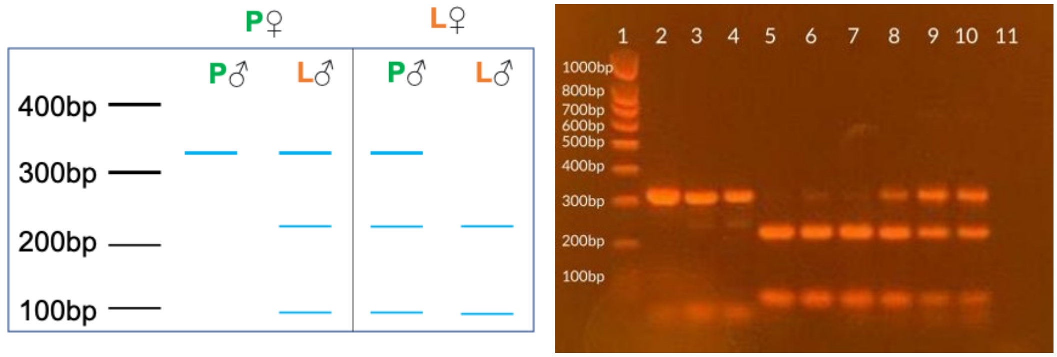

***Supplement Figure 2:*** *Sanger Sequencing of P, L, and F₁s confirms SNP_A1. Sanger Sequencing results were mapped to a read from a lecithotrophic worm. SNP_A1 is highlighted in pink. Cut sequence for HinFI is shown. Row image taken from SnapGene.*

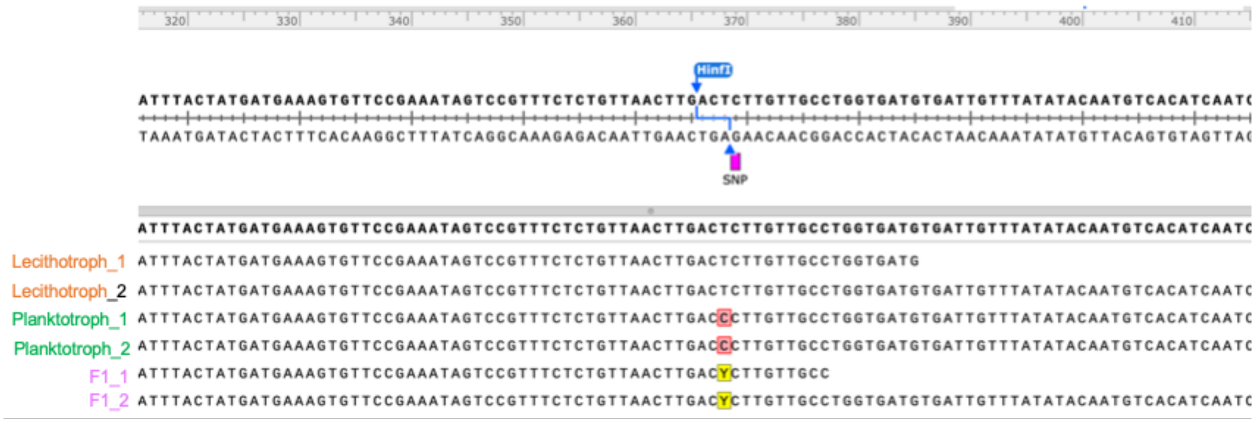

**Supplemental Methods**

We designed primers to amplify a 321bp region around SNP_A1. (Primer A3. forward: CATACAGTTTGTGAATATGC and reverse: TGTGAGGTCATCCATGTTGG in the reverse direction. We started with a longer 516bp primer set (A1) but due to heterozygosity in this region of the genome we found indels were occasionally affecting our PCR band length so we switched to the shorter PCR product after Cross8.) We used manufacturer's recommendations for PCR at 56°C annealing temperature (NEB Q5 Hot Start High Fidelity DNA Polymerase). PCR products were incubated with the HinFI enzyme as recommended and checked via gel electrophoresis. When cut, the bands of the PCR product are 225 and 96 base pairs, which is easily visualized (SFig. 1). Therefore, we assigned paternity based on the number of bands. We used Sanger sequencing to confirm the SNP pattern in our PCR product using two known P, L and $F_1$ individuals in both the forward and reverse direction (SFig. 2).

**Supplemental Results**

*Supplement Table 1. Outcome of paternity for each cross. Females chose to mate with like-males 83% of the time. There is only one instance out of the 19, where both females chose the opposite male (gray). Four crosses where a L female chose a P male are orange. One cross where a P female chose a L male is green. *Three crosses with a heterozygote parent. [+]One cross with a possible heterozygote parent.*

| | Name | Female | Male | Direction | Female Order | Bands |
|---|---|---|---|---|---|---|
| 1 | Cross 5 | P | P | Same | NA | 1 |
| | | L | L | Same | | 2 |
| 2 | Cross 6 | P | P | Same | P | 1 |
| | | L | L | Same | | 2 |
| 3 | Cross 9 | P | P | Same | P | 1 |
| | | L | L | Same | | 2 |
| 4 | Cross 12 | P | P | Same | P | 1 |
| | | L | L | Same | | 2 |
| 5 | Cross 15 | P | P | Same | L | 1 |
| | | L | L | Same | | 2 |
| 6 | Cross 18 | P | P | Same | L | 1 |
| | | L | L | Same | | 2 |
| 7 | Cross 22 | P | P | Same | P | 1 |
| | | L | L | Same | | 2 |
| 8 | Cross 23 | P | P | Same | L | 1 |
| | | L | L | Same | | 2 |
| 9 | Cross 26 | P | P | Same | L | 1 |
| | | L | L | Same | | 2 |
| 10 | Cross 27* | P | P | Same | P | 1 |

| | | | | | | |
|---|---|---|---|---|---|---|
| | | L | L | Same | | 2:3 |
| 11 | Cross 30* | P | P | Same | L | 1 |
| | | L | L | Same | | 2:3 |
| 12 | Cross 31 | P | P | Same | P | 1 |
| | | L | L | Same | | 2 |
| 13 | Cross 34* | P | P | Same | L | 1 |
| | | L | L | Same | | 2:3 |
| 14 | Cross 11 | P | P | Same | L | 1 |
| | | L | P | Opposite (L) | | 3 |
| 15 | Cross 17 | P | P | Same | L | 1 |
| | | L | P | Opposite (L) | | 3 |
| 16 | Cross 19 | P | P | Same | L | 1 |
| | | L | P | Opposite (L) | | 3 |
| 17 | Cross 29 | P | P | Same | L | 1 |
| | | L | P | Opposite (L) | | 3 |
| 18 | Cross 33+ | P | L | Opposite (P) | P | 3 |
| | | L | L | Same | | 2 |
| 19 | Cross 24 | P | L | Opposite (P) | P | 3 |
| | | L | P | Opposite (L) | | 3 |
| 20 | Cross 2 | L | L | Same | | 2 |
| 21 | Cross 3 | P | P | Same | | 1 |
| 22 | Cross 4 | P | P | Same | | 1 |
| 23 | Cross7 | L | L | Same | | 2 |
| 24 | Cross 8 | L | L | Same | | 2 |
| 25 | Cross 10 | L | L | Same | | 2 |
| 26 | Cross 13 | P | P | Same | | 1 |
| 27 | Cross 16 | L | L | Same | | 2 |
| 28 | Cross 20 | P | P | Same | | 1 |
| 29 | Cross 21 | P | P | Same | | 1 |
| 30 | Cross 25 | L | L | Same | | 2 |
| 31 | Cross 28 | P | P | Same | | 1 |

| | | | | | | |
|---|---|---|---|---|---|---|
| 32 | Cross 35 | L | L | Same | | 2 |
| 33 | Cross 1 | L | P | Opposite (L) | | 3 |
| 34 | Cross 14 | L | P | Opposite (L) | | 3 |

In the three crosses where heterozygosity was indicated, we saw a pattern of 50:50 for 2:3 band offspring(SFig. 1). In all cases were able to assign paternity in these crosses to the L male. There are two possible scenarios from this result: 1) A heterozygous L father would produce 50% offspring with the LL genotype. 2) A heterozygous L mother would also produce the same 2:3 band pattern, but only if she mated with the LL male. (If she mated with a PP male the bands would be 1:2, and if both parents were heterozygotes we would expect a 1:2:1 ratio of bands). Either scenario does not change the assignment of paternity to the L father.

We considered the possibility that a P individual might be heterozygous in our crosses, which also would not affect our results. In this scenario, the pooled offspring from a P mother would have three bands, and would falsely indicate an opposite-type mating. There are only two crosses from a P mother that resulted in three bands: Cross24 and Cross33. Both males in Cross24 were genotyped and they conformed to expected homozygous genotypes. So the only cross that could possibly have a false assignment due to unexpected heterozygosity is Cross33, where it appears that the PP females mated with the LL male. When we see three bands on the gel for P mothers, there are three possible options: 1) PP female mated with LL male, 2) PP female mated with LP Male or 3) PL female mated with either male. Scenario one and two give the same paternity of a L father, assuming it is again the L male that is heterozygous. We think it is unlikely that the P female is a heterozygote in this scenario as no females and no P individuals have been genotyped as heterozygotes. It is more likely that both P and L females chose to mate with the L male. We cannot be absolutely sure of the parental genotypes in this cross, but removing it from the analysis does not change the overall result.
